# Multivariate genome-wide association study identifies 780 unique genetic loci associated with cortical morphology

**DOI:** 10.1101/2020.10.22.350298

**Authors:** Alexey A. Shadrin, Tobias Kaufmann, Dennis van der Meer, Clare E. Palmer, Carolina Makowski, Robert Loughnan, Terry L. Jernigan, Tyler M. Seibert, Donald J Hagler, Olav B. Smeland, Yunhan Chu, Aihua Lin, Weiqiu Cheng, Guy Hindley, Wesley K. Thompson, Chun C. Fan, Dominic Holland, Lars T. Westlye, Oleksandr Frei, Ole A. Andreassen, Anders M. Dale

## Abstract

Brain morphology has been shown to be highly heritable, yet only a small portion of the heritability is explained by the genetic variants discovered so far. Here we exploit the distributed nature of genetic effects across the brain and apply the Multivariate Omnibus Statistical Test (MOSTest) to genome-wide association studies (GWAS) of vertex-wise structural magnetic resonance imaging (MRI) cortical measures from N=35,657 participants in the UK Biobank. We identified 695 loci for cortical surface area and 539 for cortical thickness, in total 780 unique genetic loci associated with cortical morphology. This reflects an approximate 10-fold increase compared to the commonly applied univariate GWAS methods. Power analysis indicates that applying MOSTest to vertex-wise structural MRI data triples the effective sample size compared to conventional univariate GWAS approaches. Functional follow up including gene-based analyses implicate 10% of all protein-coding genes and point towards pathways involved in neurogenesis and cell differentiation.

## Introduction

Variability in brain morphology is highly heritable, with twin studies estimating heritability for global measures at 89% for total surface area and 81% for mean cortical thickness^1^ and regional measures (adjusting for whole brain measures) at up to 46% for cortical area and 57% for thickness^2^. GWAS is a powerful tool for identifying genetic variants that shape the human cortex, but the full breadth of reported heritability estimates has yet to be uncovered. The most recent large-scale GWAS of brain MRI data (N=51,665) from the ENIGMA consortium identified 187 and 50 loci associated with global and regional cortical surface area and thickness, respectively^3^. The relatively low yield despite high heritabilities of brain morphology is likely due to high polygenicity and small effect size (discoverability) per locus^4^.

Both imaging genetics^4^ and gene expression studies^5^ suggest that genetic effects are distributed across cortical regions, such that variants influencing one cortical region are also likely to affect other cortical regions. Multivariate statistical methods are naturally tailored to model distributed and pleiotropic genetic effects. We recently developed a Multivariate Omnibus Statistical Test (MOSTest)^6^ that aggregates effects across spatially distributed phenotypes, such as cortical thickness, boosting our ability to detect variant-phenotype associations. We showed that applying MOSTest to cortical morphology region of interest (ROI) measures in the UK Biobank substantially increased loci discovery^6^ compared to the commonly applied approach used by the ENIGMA consortium^3^, here referred to as the min-P approach. For each genetic variant tested for association with multiple phenotypes, min-P considers only the most significant p-value and corrects it for the effective number of phenotypes analyzed, thus failing to exploit shared genetic architecture across brain regions. In contrast, MOSTest relies on the distributed nature of genetic influences across brain regions and allows detection of genetic variants with weak effects in multiple brain regions. We have shown that the discoverability of GWAS variants underlying regional cortical area and thickness depends on the specific parcellation of cortical regions used, and that parcellations based on genetic correlations from twin studies perform better than genetically un-informed schemes^4^. Here we show that the combined genetic yield (number of loci discovered) for cortical area and thickness can be boosted when using MOSTest (yielding a 3.8-fold increase relative to min-P), and boosted further when moving from a region-based approach to a more fine-grained vertex-wise approach (additional 1.8-fold increase). Uncovering the detailed genetic architecture of cortical area and thickness will provide insight into the underlying neurobiology of the human brain, and give a better understanding of brain-related human traits, such as cognition^7^, as well as neurological^8^ and psychiatric diseases^9^.

## Results

### Genetic loci discovery

Using MOSTest^6^, we performed a multivariate GWAS of cortical morphology, such that the significance of each locus was estimated after aggregating its effects across all vertices (1284 data points each for thickness and area). This was conducted separately for cortical surface area and thickness in 35,657 individuals from UK Biobank. Cortical morphology estimates were residualized for modality-specific global brain measures prior to analysis in order to estimate regional cortical effects relative to global brain measures. Measurements from left and right hemispheres were included separately (not averaged). We identified 695 and 539 loci, respectively, equating to 780 unique loci associated with cortical morphology. Prior to performing the MOSTest analysis, individual cortical area and thickness measures were residualized for age, sex, scanner site, proxy of surface reconstruction quality, the first twenty genetic principal components, and a participant-specific global measure (either total area or average thickness). Measurements from left and right hemispheres were not merged. For comparison, we repeated this procedure aggregating over 68 ROIs from the Desikan-Killiany parcellation. This resulted in the discovery of 370 loci for cortical surface area and 181 loci for cortical thickness, such that the vertex-wise MOSTest analysis provided a 1.9-fold and 3.0-fold increase in yield over the region-based MOSTest analysis, respectively. Applying the min-P approach to Desikan-Killiany ROIs resulted in further reduction in the number of loci discovered (88 for cortical surface area; 44 for cortical thickness). This represents a 4.2-fold and 4.1-fold decrease compared to the MOSTest ROI-based analysis, and a 7.9-fold and 12.3-fold decrease compared to vertex-wise MOSTest analysis, respectively. Manhattan plots are presented in Fig. 1, with corresponding QQ plots in Supplementary Fig. 1. Numbers of loci discovered with different approaches are shown in Supplementary Table 1. Specific loci discovered in each analysis are listed in Supplementary Tables 2 - 7.

**Fig. 1:**
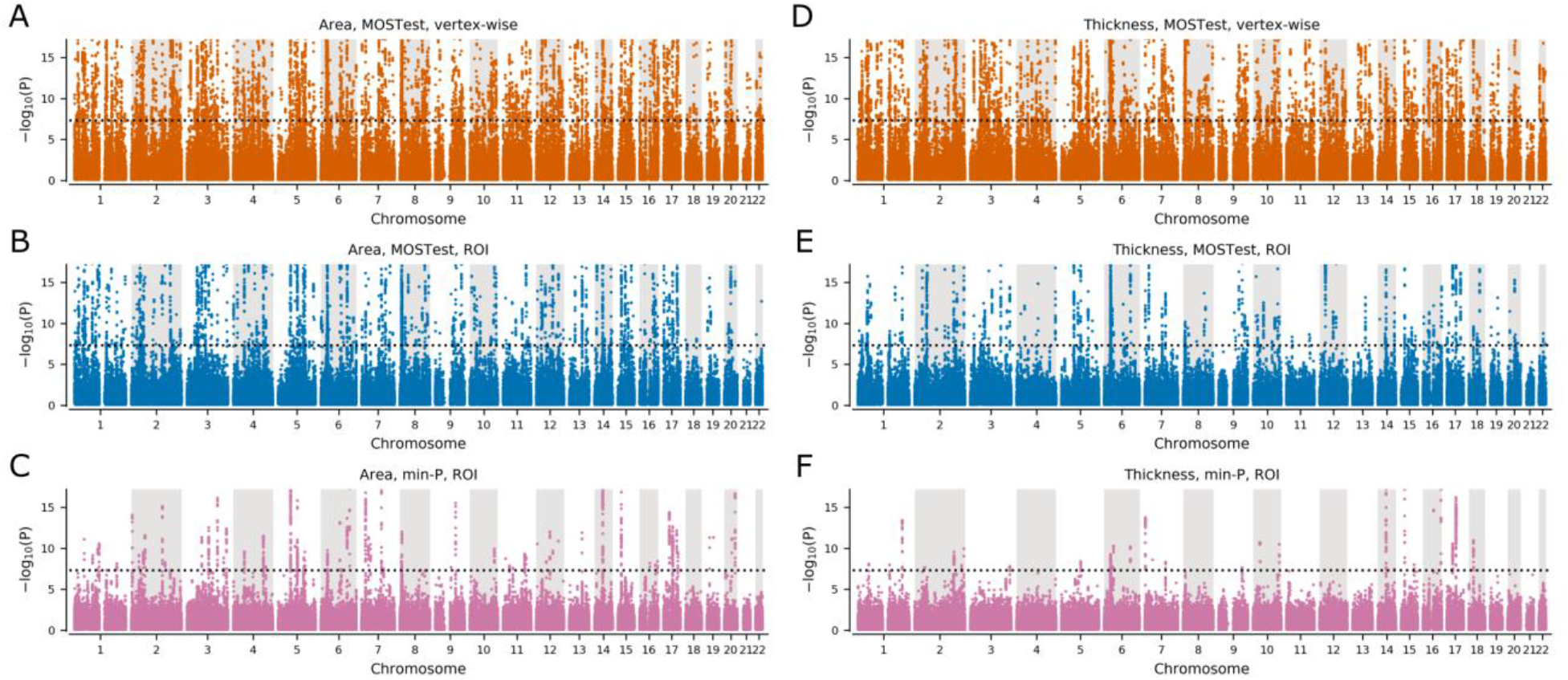
Manhattan plots for cortical surface area and cortical thickness. (A) Area, MOSTest, vertex-wise: N=695 loci. (B) Area, MOSTest, ROI: N=370 loci. (C) Area, min-P, ROI: N=88 loci. (D) Thickness, MOSTest, VW: N=539 loci. (E) Thickness, MOSTest, ROI: N=181 loci. (F) Thickness, min-P, ROI: N=44 loci. Black dotted horizontal lines show genome-wide significance threshold (P=5E-8). Loci; independent genome-wide significant (P<5E-8). Y-axes are truncated at -log10(P)=17.2 to highlight the region around genome-wide significance threshold. ROI = region of interest.

To compare the vertex-wise MOSTest results with the most recent ENIGMA GWAS^3^, we also applied the ENIGMA-based definition of genetic locus. This resulted in 1598 and 1054 unique loci for cortical area and thickness respectively, and a total of 1735 unique loci for cortical morphology identified in the vertex-wise MOSTest analysis (Supplementary Tables 8 - 9).

### Power analysis

To estimate the proportion of additive genetic variance explained by genome-wide significant SNPs identified by either MOSTest or min-P as a function of sample size, we used the MiXeR tool^10^ (Fig. 2). The horizontal shift of the curve indicates that the effective sample size of MOSTest is around threefold that of min-P. We estimate that with the current UK Biobank sample (N=35,657), 11.6% and 7.0% of the additive genetic variance in cortical surface area and thickness, respectively, can be explained by genome-wide significant loci from the vertex-wise MOSTest analysis. (Fig. 2). In contrast, the min-P approach identifies 1.3% and 0.2% of the explained additive genetic variance for area and thickness, respectively (Fig. 2). The power-analysis indicates that 32.2% and 24.0% of the additive genetic variance in cortical surface area and thickness, respectively, will be discovered in the full UK Biobank sample of N=100,000 using the MOSTest vertex-wise approach (Fig. 2). Further, the proportion of explained variance with the min-P approach in the full UK Biobank sample is estimated to be lower than the yield of MOSTest in the present sample size (Fig. 2).

**Fig. 2:**
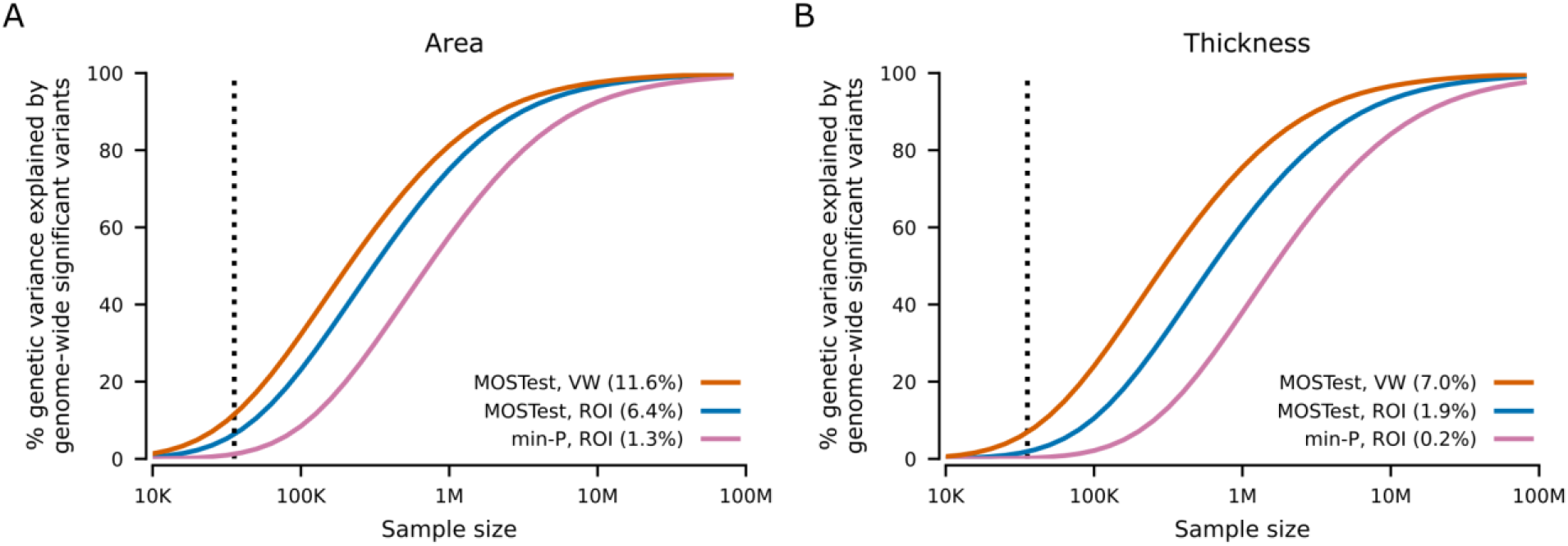
Estimated percent of additive genetic variance explained by genome-wide significant SNPs as a function of sample size. Percentages of genetic variance explained by identified SNPs (p<5E-8) from multivariate GWAS (MOSTest VW) of area (A) and thickness (B) with current sample size (N=35,657, vertical dotted line) are shown in parentheses, with MOSTest ROI and min-P ROI for comparison. VW = vertex-wise. ROI = region of interest.

### Gene-level analysis

Through gene-level analyses of the vertex-wise MOSTest GWAS using MAGMA^11^, we found that 1647 and 1412 genes, out of a total of 19036 protein-coding genes, were significantly associated with area and thickness, respectively (Supplementary Table 10). We also performed competitive gene-set analyses restricted to the Gene Ontology biological processes category (containing 7343 pathways). This resulted in 204 and 184 significant (p<0.05/7343) gene sets associated with area and thickness, respectively. The most significantly associated pathways were related to neuronal development and cell differentiation, with the top 10 shown in Fig. 3.

**Fig. 3:**
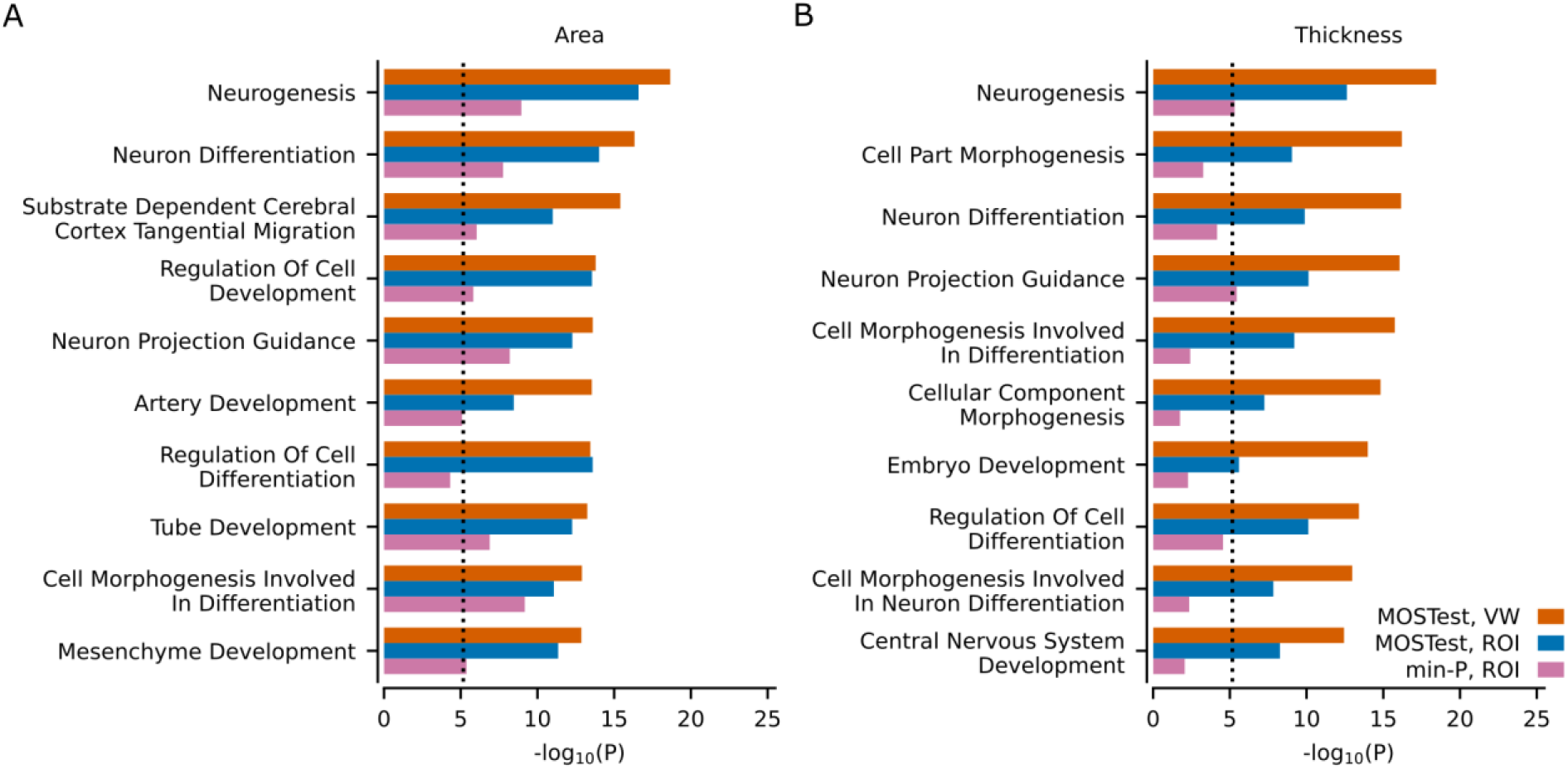
Gene-set analyses with MAGMA. Results from the gene-set analysis based on multivariate GWAS on area (A) and thickness (B). Ten most significant Gene Ontology sets (N=7343) in the MOSTest VW analysis are listed on the y-axis, in comparison with MOSTest ROI and min-P ROI. Corresponding uncorrected -log10(p-values) are shown on the x-axis. P-values were obtained using MAGMA analysis as implemented in FUMA. Vertical dotted line shows Bonferroni correction threshold (p=0.05/7343). VW = vertex-wise. ROI = region of interest.

For comparison, we also performed the same analyses on the ROI-based MOSTest and min-P GWAS summary statistics, resulting in 198 area and 66 thickness gene sets for ROI-based MOSTest and 60 area and 4 thickness gene sets for min-P. As shown in Supplementary Figs. 2 and 3, the vertex-wise MOSTest approach led to much greater significance for nearly all pathways identified. Interestingly, the most significant pathways identified by vertex-wise MOSTest are tightly connected with critical neurobiological processes implicated in brain development while top findings in the min-P analysis are less specific. The distributed effects of identified variants across different brain regions are also illustrated by brain maps, highlighting the mixture of effects across the cortex (Supplementary Fig. 4).

## Discussion

We identified 695 loci for cortical surface area and 539 for cortical thickness, in total 780 unique genetic loci associated with cortical morphology. This reflects an approximate 10-fold increase compared to the commonly applied univariate GWAS methods. Our study highlights the greatly improved yield obtained with the multivariate method compared to conventional univariate GWAS approach, which stems from the multivariate nature of brain morphology phenotypes, representing continuous maps per individual. The present results support the hypothesis that the genetic determinants of variability in brain morphology are extensively shared across multiple regions^6^. Our findings further underscore the complex molecular mechanisms shaping the human brain, which we show are largely related to neurodevelopmental processes.

Twin studies have suggested the largely independent nature of cortical surface area and thickness^1^. The genetic correlation between them estimated using linkage disequilibrium score regression (LDSR) is rg=-0.32 (p=6.5E-12)^3^. Here we identify the specific loci involved and show that these cortical phenotypes share a large proportion of genomic loci. Out of a total of 695 loci for cortical area and 539 loci for cortical thickness, 454 loci (58.2% of the total number of unique loci) were overlapping. These findings illustrate how measures of genetic correlation fail to fully capture the extent to which the genetic influences of two phenotypes are interrelated. LDSR and twin analyses depend on the consistency of effect directions across phenotypes. In contrast, the analysis performed here consider non-null loci as overlapping if they are both significant and in linkage disequilibrium, regardless of effect directions. Overlapping genetic architecture across brain regions despite the absence of strong genetic correlations are therefore plausible due to common molecular toolkits involved in neurodevelopment across brain regions^12^. This is in line with Allen Brain Atlas maps of the adult human brain^13^, showing regions with high similarity in gene expression between cortical structures consistent with the notion that the basic architecture across the entire cortex is similar or “canonical”^14^. This may also explain the shared genetic architecture observed for many brain-related traits and disorders^15-17^. Accounting for the distributed signal across the cortex in a multivariate framework allowed us to boost power for discovery compared to traditional univariate approaches, such as min-P.

Our gene-level analyses indicated that, with the current sample size, 10% of all protein-coding genes were significantly associated with brain morphology (either cortical area or thickness). Gene-set analyses for both area and thickness confirmed involvement of pathways recently reported by ENIGMA^3^, but with greater statistical significance. We additionally found strong evidence for the involvement of several genetic pathways regulating neuronal development and differentiation that were not identified by the min-P approach, implicating key biological processes regulating human surface area expansion and increases in thickness. This also corroborates the strong statistical signals and suggests that we are capturing true biological mechanisms that were missed by previous methodologies. These novel findings of neurobiological underpinnings associated with brain morphology provide a framework for follow-up experimental studies to identify the complex polygenic mechanisms involved in human brain development^18^. Further, the findings implicating neuronal development and cell differentiation can facilitate experimental studies to gain better insight into the pathobiological mechanisms of brain-related diseases including psychiatric disorders^19^, where we need to understand the role of polygenic mechanisms^20^.

Compared to the current largest brain morphology GWAS (N=50K)^3^, analyzing parcellation-free, vertex-wise data with MOSTest increased the yield of significant loci 8.5-fold for cortical surface area and 21.1-fold for cortical thickness, despite the lower sample size in our study (N=35K). Of note, while being generally consistent, our protocol differs in a few aspects from the previous GWAS^3^, where global measures were included in the principal analysis and data for cortical regions were averaged across right and left hemispheres. Using the Desikan-Killiany parcellation approximately 2.0 times more variants were identified for cortical surface area than for cortical thickness both with the min-P and the MOSTest (Supplementary Table 1). In contrast, there were 1.3 times more loci for area compared to thickness when using the MOSTest for parcellation-free vertex-wise data. (Supplementary Table 1). The observed difference in loci yield may be due to differing degrees of mismatch between parcellation schemes and actual architecture of the phenotypes. This seems to be particularly relevant for thickness, where variant effects obtained from an ROI parcellation scheme may be underestimated compared to the vertex-wise approach. This result may explain why parcellation schemes better reflecting the genetic architecture of the cortex improve detectability in imaging genetics studies^4^.

The boost in statistical power using the multivariate vertex-wise approach is equivalent to a more than three-fold increase in effective sample size for both area and thickness (Fig. 2). Our analysis suggests that the substantial gain in power provided by MOSTest is projected to explain approximately 32.2% and 24.0% of the additive genetic variance for cortical surface area and thickness, respectively, upon completion of UK Biobank’s target neuroimaging sample (N=100,000)^21^ (Fig. 2). It is possible that multivariate approaches will also boost discovery of genetic associations with other human phenotypes that exhibit shared signal between traits.

To conclude, we have identified 780 unique loci associated with human brain morphology, highlighting its polygenic nature and providing the foundation for functional follow-up experiments. While this study is focused solely on UK Biobank, the generalizability and flexibility of this approach allows its incorporation into large-scale meta-analyses like ENIGMA^22^, offering unique opportunities for major advances in our understanding of the genetic determinants of brain morphology.

## Materials and Methods

### Sample

Genotypes, MRI scans, demographic and clinical data were obtained from the UK Biobank under accession number 27412. For this study, we selected white British individuals (as derived from both self-declared ethnicity and principal component analysis^23^) who had undergone the neuroimaging protocol. The resulting sample contained 35,657 individuals with a mean age of 64.4 years (standard deviation 7.5 years), 51.7% female.

### Data processing

T1-weighted structural MRI scans were processed with the FreeSurfer v5.3 standard “recon-all” processing pipeline^24^ to generate 1284 non-smoothed vertex-wise measures (ico3 downsampling with the medial wall removed) and 68 ROI measures (based on the Desikan-Killiany parcellation) summarizing cortical surface area and thickness. All measures were pre-residualized for age, sex, scanner site, a proxy of surface reconstruction quality (FreeSurfer’s Euler number^25^), the first twenty genetic principal components, and a global measure specific to each set of variables: total cortical surface area and mean cortical thickness for the regional area and thickness measurements correspondingly. Subsequently, a rank-based inverse normal transformation was applied to the residualized measures. We used UK Biobank v3 imputed genotype data^23^, carrying out standard quality-checks as described previously^6^, and setting a minor allele frequency threshold of 0.5%, leaving 9 million variants. Variants were tested for association with cortical surface area and cortical thickness at each vertex and each ROI separately using the standard univariate GWAS procedure. Resulting univariate p-values and effect sizes were further combined in the MOSTest and min-P analyses to identify area- and thickness-associated loci.

### MOSTest analysis

Consider *N* variants and *M* (pre-residualized) phenotypes. Let *z*_*ij*_ be a z-score from the univariate association test between i^th^ variant and j^th^ (residualized) phenotype and *z*_*i*_ = (*z*_*i*1_, … ,*z*_*iM*_) be the vector of z-scores of the i^th^ variant across *M* phenotypes. Let *Z* = {*z*_*ij*_} be the matrix of z-scores with variants in rows and phenotypes in columns. For each variant consider a random permutation of its genotypes and let 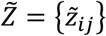 be the matrix of z-scores from the univariate association testing between variants with permuted genotypes and phenotypes. A random permutation of genotypes is done once for each variant and the resulting permuted genotype is tested for association with all phenotypes, therefore preserving correlation structure between phenotypes.

Let 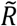 be the correlation matrix of 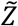, and 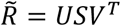 is its singular valued decomposition (*U* and *V* – orthogonal matrixes, *S*– diagonal matrix with singular values of 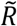 on the diagonal). Consider the regularized version of the correlation matrix 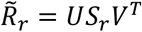, where *S*_*r*_ is obtained from *S* by keeping *r* largest singular values and replacing the remaining with *r*_th_largest. The MOSTest statistics for i^th^ variant (scalar) is then estimated as 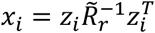, where regularization parameter *r* is selected separately for cortical area and thickness to maximize the yield of genome-wide significant loci. In this study we observed the largest yield for cortical surface area with *r*=10; the optimal choice for cortical thickness was *r*=20 (Supplementary Fig. 5). The distribution of the test statistics under null 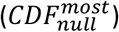 is approximated from the observed distribution of the test statistics with permuted genotypes, using the empirical distribution in the 99.99 percentile and Gamma distribution in the upper tail, where shape and scale parameters of Gamma distribution are fitted to the observed data. The p-value of the MOSTest test statistic for the i^th^ variant is then obtained as 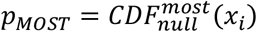.

### min-P analysis

Similar to the MOSTest analysis, consider *N* variants and *M* preresidualized phenotypes. Let *z*_*ij*_ be a z-score from the univariate association test between i^th^ variant and j^th^ (residualized) phenotype and *z*_*i*_ = (*z*_*i*1_, … ,*z*_*iM*_) be the vector of z-scores of the i^th^ variant across *M* phenotypes. The min-P statistics for the i^th^ variant is then estimated as 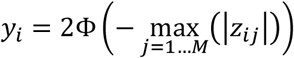, where Φ is a cumulative distribution function of the standard normal distribution. The distribution of the min-P test statistics under null 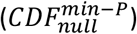 is approximated from the observed distribution of the test statistics with permuted genotypes, using the empirical distribution in the 99.99^th^ percentile and Beta distribution in the upper tail, where shape parameters of Beta distribution (*α* and *β*) are fitted to the observed data. The p-value of the min-P test statistic for the i^th^ variant is then obtained as 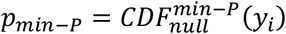.

### Locus definition

Genetic loci were defined based on association summary statistics produced with MOSTest and min-P following the protocol implemented in FUMA^26^ with default parameters. The protocol can be summarized as the following:

1. Independent significant genetic variants are identified as variants with p-value<5E-8 and linkage disequilibrium (LD) r2<0.6 with each other.
2. A subset of these independent significant variants with LD r2<0.1 are then selected as lead variants.
3. For each independent significant variant all candidate variants are identified as variants with LD r2≥0.6 with the independent significant variant.
4. For a given lead variant the borders of the genomic locus are defined as min/max positional coordinates over all corresponding candidate variants.
5. Loci are then merged if they are separated by less than 250kb.

Alternatively, to facilitate comparison with the current largest brain morphology GWAS^3^, we also counted genetic loci applying locus definition similar to that used by ENIGMA. Briefly, the association summary statistics produced with either MOSTest or min-P were clumped with PLINK^27^ using p-value threshold of 5E-8 (--clump-p1) and linkage disequilibrium cutoffs of 1 Mb (--clump-kb) and r2 < 0.2 (--clump-r2). Obtained clumps of variants were considered as independent genome-wide significant genetic loci.

### MiXeR analysis

MOSTest and min-P p-values were analyzed with the MiXeR tool^10^ to estimate the proportion of additive genetic variance explained by genome-wide significant SNPs as a function of sample size. Right censoring (MiXeR option: --z1max 5.45) was applied to mitigate extreme effects which may lead to biased estimates.

### Gene-level analysis

We carried out MAGMA-based gene analyses using default settings, which entail the application of a SNP-wide mean model to GWAS summary statistics, with the use of the 1000 Genomes Phase 3 EUR reference panel. Gene-set analyses were done in a similar manner, restricting the sets under investigation to those that are part of the Gene Ontology biological processes subset (N=7343), as listed in the Molecular Signatures Database (MsigdB) v7.0.

## Data availability

The data incorporated in this work were gathered from the public UK Biobank resource.

## Code availability

MOSTest code is publicly available at https://github.com/precimed/mostest (GPLv3 license).

## Acknowledgements

We were funded by the Research Council of Norway (276082, 213837, 223273, 204966/F20, 229129, 249795/F20, 225989, 248778, 249795), the South-Eastern Norway Regional Health Authority (2013-123, 2014-097, 2015-073, 2016-064, 2017-004), Stiftelsen Kristian Gerhard Jebsen (SKGJ-Med-008), the EEA Grant 2014-2021 under the project contract No 6/2019, the European Research Council (ERC) under the European Union’s Horizon 2020 research and innovation programme (ERC Starting Grant, Grant Agreement No. 802998) and National Institutes of Health (R01MH100351, R01GM104400, NIDA/NCI: U24DA041123). This work was partly performed on the TSD (Tjeneste for Sensitive Data) facilities, owned by the University of Oslo, operated and developed by the TSD service group at the University of Oslo, IT-Department (USIT). Computations were also performed on resources provided by UNINETT Sigma2—the National Infrastructure for High Performance Computing and Data Storage in Norway. This work used the Extreme Science and Engineering Discovery Environment (XSEDE) including COMET and OASYS resources at the UCSD through allocation TG-IBN200001.

