## Supplementary Figures 1 - 5 for "Multivariate genome-wide association study identifies 780 unique genetic loci associated with cortical morphology"

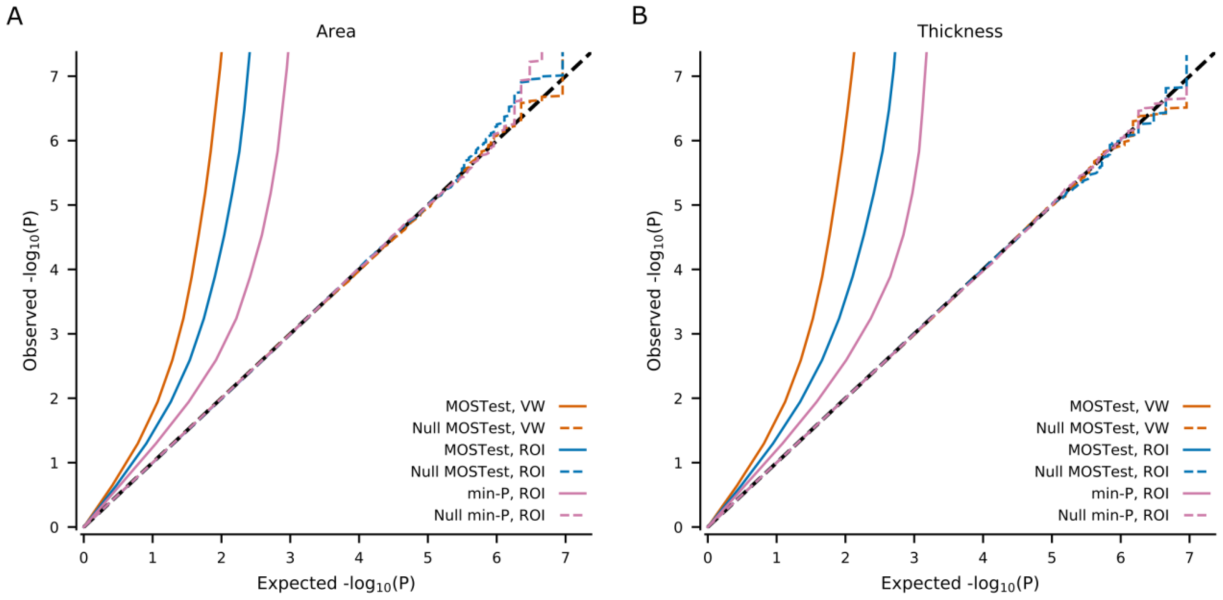


**Supplementary Fig. 1: QQ plots.**

QQ plots show summary statistics for MOSTest VW, MOSTest ROI and min-P ROI for area (A) and thickness (B) and their null (permuted) versions. VW = vertex-wise. ROI = region of interest.

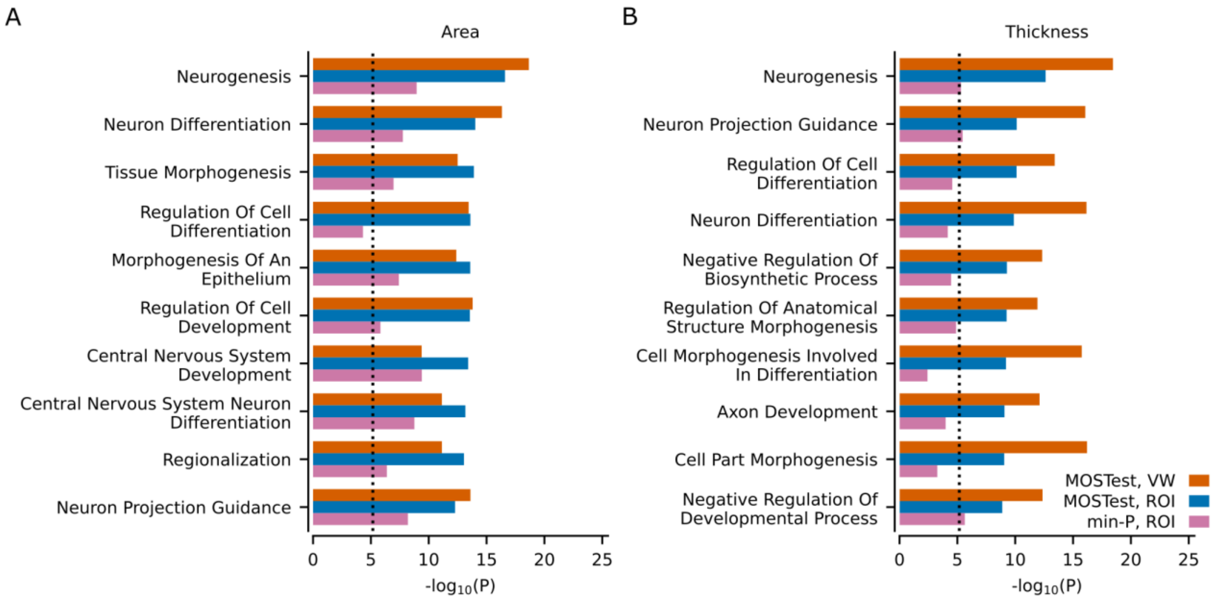


**Supplementary Fig. 2: Gene-set analyses with MAGMA prioritized based on MOSTest ROI results.**

Results from the gene-set analysis based on multivariate GWAS on area and thickness. Ten most significant Gene Ontology sets (N=7343) in the MOSTest ROI analysis are listed on the y-axis for (A) Area (B) Thickness. Corresponding uncorrected -log10(p-values) are shown on the x-axis. P-values were obtained using MAGMA analysis as implemented in FUMA. Vertical dotted line shows Bonferroni correction threshold (p=0.05/7343). VW = vertex-wise. ROI = region of interest.

6

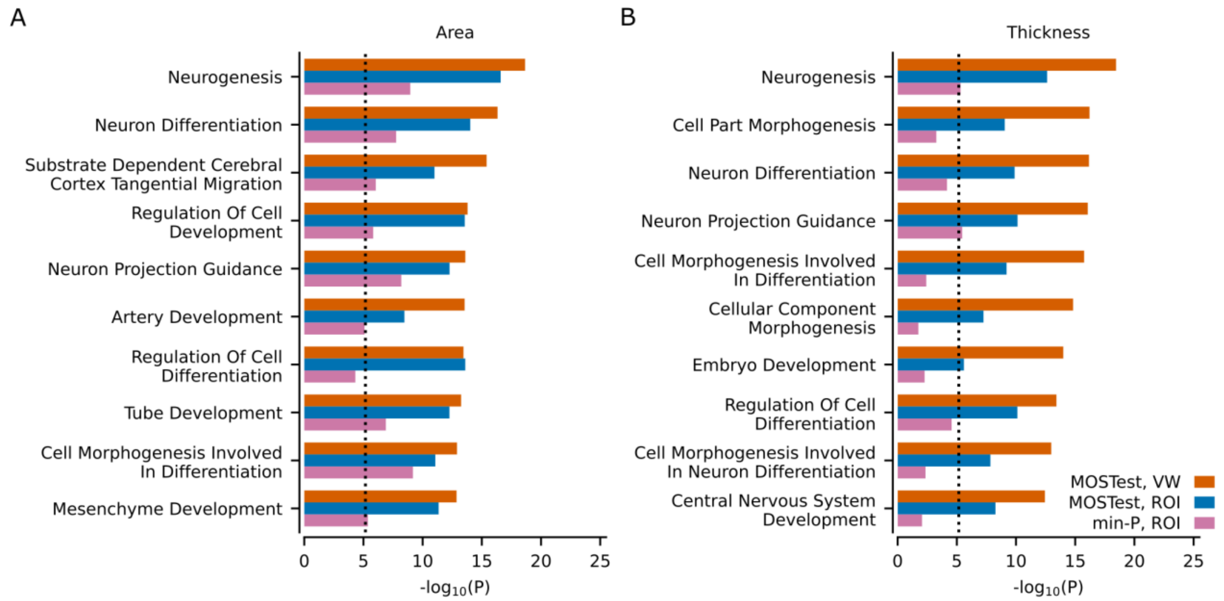


**Supplementary Fig. 3: Gene-set analyses with MAGMA prioritized based on min-P ROI results.**

Results from the gene-set analysis based on multivariate GWAS on area and thickness. Ten most significant Gene Ontology sets (N=7343) in the min-P ROI analysis are listed on the y-axis for (A) Area, (B) Thickness. Corresponding uncorrected -log10(p-values) are shown on the x-axis. P-values were obtained using MAGMA analysis as implemented in FUMA. Vertical dotted line shows Bonferroni correction threshold (p=0.05/7343). VW = vertex-wise. ROI = region of interest.

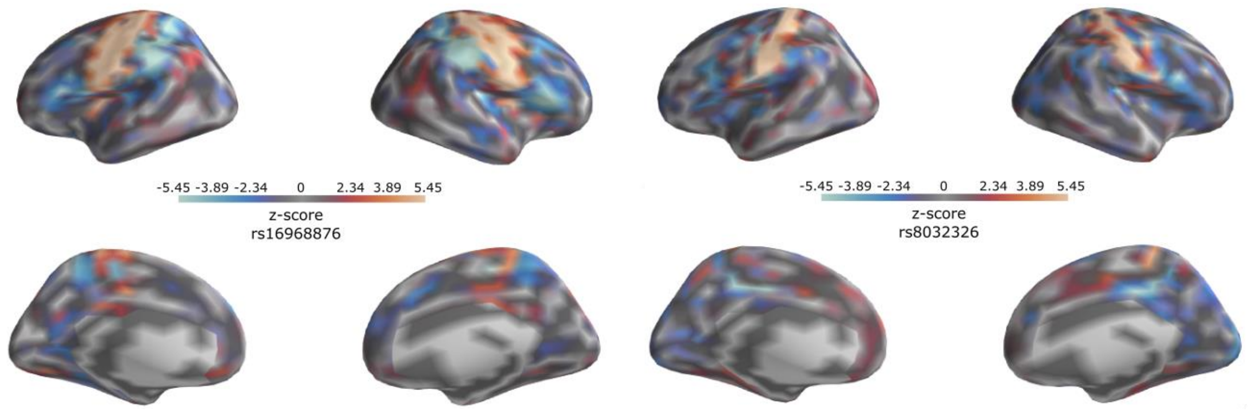


**Supplementary Fig. 4: Z-scores from the univariate GWAS for the top genetic loci (ENIGMA-based locus definition) identified in the vertex-wise MOSTest analysis.**

(left): top lead variant (rs16968876, chr15:39636877) associated with cortical surface area.

(right): top lead variant (rs8032326, chr15:39619456) associated with cortical thickness.

9

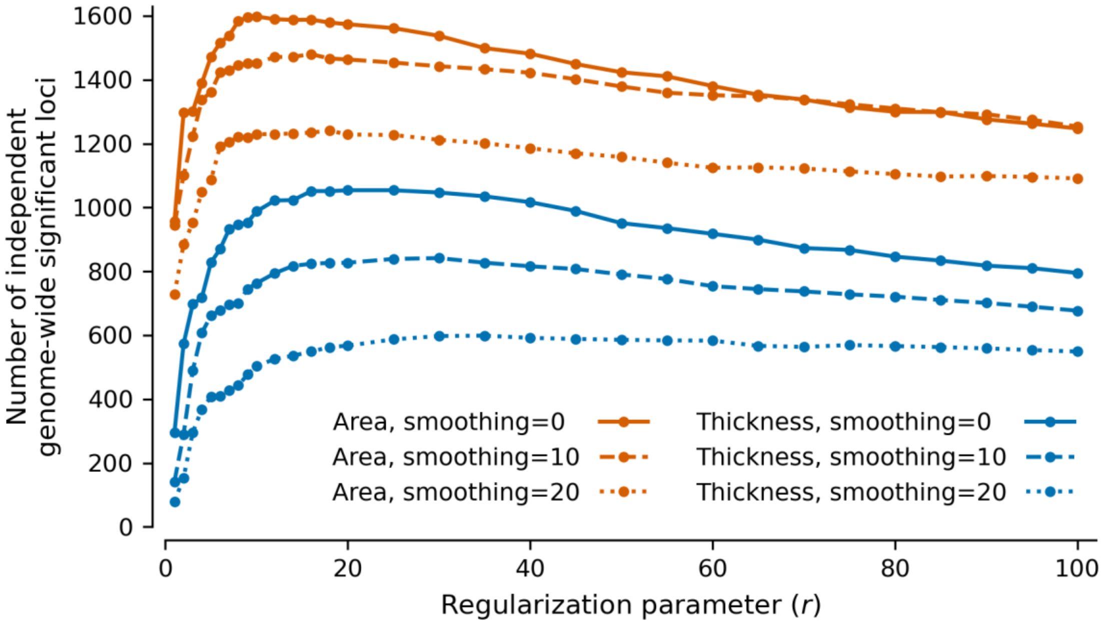


**Supplementary Fig. 5**: **Effect of regularization parameter on loci yield.**

Number of genome-wide significant loci (ENIGMA-based locus definition) identified for cortical area and thickness applying MOSTest with different regularization parameters ($r$=1,2,3, …, 100) to vertex-wise measures (ico3 downsampling) with various degrees of smoothing (FWHM=0, 10, 20). Smoothing was performed before downsampling to ico3.
